# Enhancing predictions of protein stability changes induced by single mutations using MSA-based Language Models

**DOI:** 10.1101/2024.04.11.589002

**Authors:** Francesca Cuturello, Marco Celoria, Alessio Ansuini, Alberto Cazzaniga

## Abstract

Protein Language Models offer a new perspective for addressing challenges in structural biology, while relying solely on sequence information. Recent studies have investigated their effectiveness in forecasting shifts in thermodynamic stability caused by single amino acid mutations, a task known for its complexity due to the sparse availability of data, constrained by experimental limitations. To tackle this problem, we introduce two key novelties: leveraging a Protein Language Model that incorporates Multiple Sequence Alignments to capture evolutionary information, and using a recently released mega-scale dataset with rigorous data pre-processing to mitigate overfitting. We ensure comprehensive comparisons by fine-tuning various pre-trained models, taking advantage of analyses such as ablation studies and baselines evaluation. Our methodology introduces a stringent policy to reduce the widespread issue of data leakage, rigorously removing sequences from the training set when they exhibit significant similarity with the test set. The MSA Transformer emerges as the most accurate among the models under investigation, given its capability to leverage co-evolution signals encoded in aligned homologous sequences. Moreover, the optimized MSA Transformer outperforms existing methods and exhibits enhanced generalization power, leading to a notable improvement in predicting changes in protein stability resulting from point mutations. Code and data are available at https://github.com/RitAreaSciencePark/PLM4Muts.

## 1 Introduction

The emergence of Protein Language Models (PLMs) marks a substantial milestone in our ability to unlock the intricate language encrypted within amino acid sequences. Leveraging transformer-based natural language processing techniques [1, 2], PLMs are instrumental in large-scale pre-training, enabling the generation of representations that are tailored for transfer learning on specific tasks. This pre-training strategy enhances the model’s ability to capture nuanced features and patterns inherent in protein sequences [3–5], thereby facilitating more effective transfer to downstream tasks such as protein structure prediction and function annotation [6–8]. Their versatility spans a wide range of applications, such as understanding protein fitness and evolutionary dynamics [9–12]. Inspired by the tight interconnection between evolutionary adaptability and structural stability, we explore the potential of these models in predicting thermodynamic stability changes induced by single amino acid mutations. The variation in stability from a wild-type protein to its mutated counterpart is determined by the difference in unfolding free energy between them. Assessing protein stability is crucial in molecular biology, given that proteins serve as fundamental components of living systems, executing diverse functions primarily dictated by their precise three-dimensional structure. Even subtle alterations in their sequence can perturb this structure, thereby impacting their function. Accurately predicting how mutations alter stability provides valuable insights into the interplay between the protein structure and function [13, 14], while enhancing our understanding of molecular evolution mechanisms [15,16]. In the context of enzyme engineering and drug development, the ability to regulate stability changes via mutations is essential for optimizing protein functionality to meet specific design requirements [17–19]. Considering that many pharmaceutical agents target proteins, understanding how non-synonymous variants influence stability facilitates the identification of promising drug targets and prediction of pharmacological treatment efficacy. Mutations in proteins can also contribute to the onset of various diseases [20–22], including cancer [23, 24]. Investigating the effects of these mutations on protein stability unveils critical insights into disease mechanisms, potentially uncovering novel therapeutic targets. The exploration of mutations’ impact on structural stability has traditionally been approached through classical methods, which rely on energy-based models or statistical potentials [25–28]. Machine learning models have also increasingly contributed, predominantly exploiting structure information [29–40], while only a few algorithms focus solely on protein sequences. [41, 42]. These techniques employ a variety of different models, ranging from Convolutional Neural Networks to Random Forest or Support Vector Machine regressions, often integrating graph-based representations of protein structures. However, the adoption of structure-based approaches can introduce a break in symmetry within the architectures, leading to a pronounced imbalance in prediction accuracy between direct and reverse mutations [43–45]. In this context, recent studies have highlighted the efficiency of transfer-learning from massively pre-trained protein language models [46–51]. Despite PLMs remarkable capabilities, the challenge of forecasting stability effects from sequence-only information is compounded by the limited availability of experimental data [52, 53], given the resource-intensive nature of experiments. The scarcity of experimental data largely influences the robustness of machine learning models, which often encounter biases arising from overlaps between training and test sets [54–56]. Addressing these evaluation issues, a recent study [45] introduces a novel test dataset specifically designed to minimize similarity with sequences present in the commonly used training sets. The assessment of state-of-the-art models reported in that work sets a benchmark for future research, highlighting the substantial impact of shared homology on the generalization capability of the models. Notably, the study reveals that all compared approaches experience drastic drops in accuracy when measured on this newly introduced test set, providing evidence of the importance of factoring out sequence similarity in predictive modeling. In response to these challenges, we fine-tune three distinct models - MSA Transformer, ESM2, and ProstT5 [57–59] - by incorporating a regression head into the architecture. Moreover, we adopt a strategic combination of a rigorous pre-processing pipeline, drastically reducing the number of training sequences, and we leverage a recently published mega-scale dataset [60]. While the mega-scale training enhances performances compared to older datasets, we also observe meaningful improvements in training with a small dataset encompassing a larger number of proteins. The results of our experiments show the superior predictive capability of the fine-tuned MSA Transformer compared to other PLMs. Overall, the optimized MSA Transformer consistently outperforms existing methods, highlighting the efficacy of incorporating information about sequence evolution to understand the effect of mutations on protein stability.

## 2 Data

### 2.1 Datasets Overview

Assessing the impact of a single mutation on protein stability using machine learning techniques is hindered by the limited amount of available experimental data, encompassing mutations associated with only a few hundred proteins. The recently released mega-scale dataset [60], generated through an innovative high-throughput experimental assay, allows the screening of thousands of single mutations, encouraging further exploration in this domain. We provide below a detailed description of the adopted datasets, labeled with the number of point mutations they contain. The initial datasets, prior to the similarity filtering outlined in Section 2.2, are obtained from a recent study [46], and are curated by including PDB-id, wild type and mutated sequences, the position and type of amino acid substitution, and the corresponding experimental measure of stability variation.

- **Test sets** :For evaluating the models, we employ the well-established *S669* dataset [45, 53] and the *ssym* dataset [44] as test sets. The *S669* dataset, consisting of 94 proteins and 669 single mutations, is deliberately designed to minimize similarity with widely used training sets, such as *S2648* [30] and VariBench [61]. On the other hand, the *ssym* dataset, encompassing 15 proteins and 342 mutations, is specifically constructed to measure prediction biases of the methods towards destabilizing mutations.
- **Small training set** :The small training set that we consider, also denoted by *S1413*, is derived from the asymmetrical *Q3421* dataset in STRUM [62] by applying the similarity filtering based on sequence alignment against the test sets (Section 2.2). It includes 1465 point mutations and comprises 97 proteins sampled with an average of 15 mutations. Moreover, we create a reverse and symmetrical counterpart of this dataset to estimate the impact of the training labels distribution asymmetry on the model learning efficiency.
- **Large training set :**The large training set that we consider, denoted by *S155329*, is generated by 1) identifying direct amino acid substitutions within the mega-scale database, initially including more than300 thousand mutations, 2) incorporating similarity filtering (Section 2.2), excluding proteins lacking any identified homologous sequence in UniClust30. The dataset comprises a total of 149 proteins and 155329 mutations. Each protein features about one thousand mutations, providing a valuable source for the uniform screening of point mutations across proteins.
- **Designed training set :**The small and large datasets display a notable disparity in the sampled mutations, despite representing the same number of proteins. This prompts an investigation into the influence of protein heterogeneity on the efficiency of the learning process. We include all of *S1413* and augment it with a randomly chosen set of 15 mutations from each protein in *S155329*, to construct a subset of the mega-scale database ensuring a mutation count comparable to that of *S1413*. With this procedure, we obtain a set of 3648 mutations, encompassing 246 proteins (*S3648*).

### 2.2 Data Preprocessing

In order to account for this effect, we employ BLASTp [63] to perform local pairwise alignments between the test and training sequences (Figure 1). A protein is removed from the training if there is an element in the test set with sequence identity exceeding 25% (relative to the alignment length), e-value below 0.01, and alignment overlap surpassing 50% (defined as the portion of the alignment length relative to the query length), following the typical cutoffs enforced in the context of remote homology detection. We ensure that the criterion for selecting training sequences is consistently met across all analyzed test sets.

**Figure 1:**
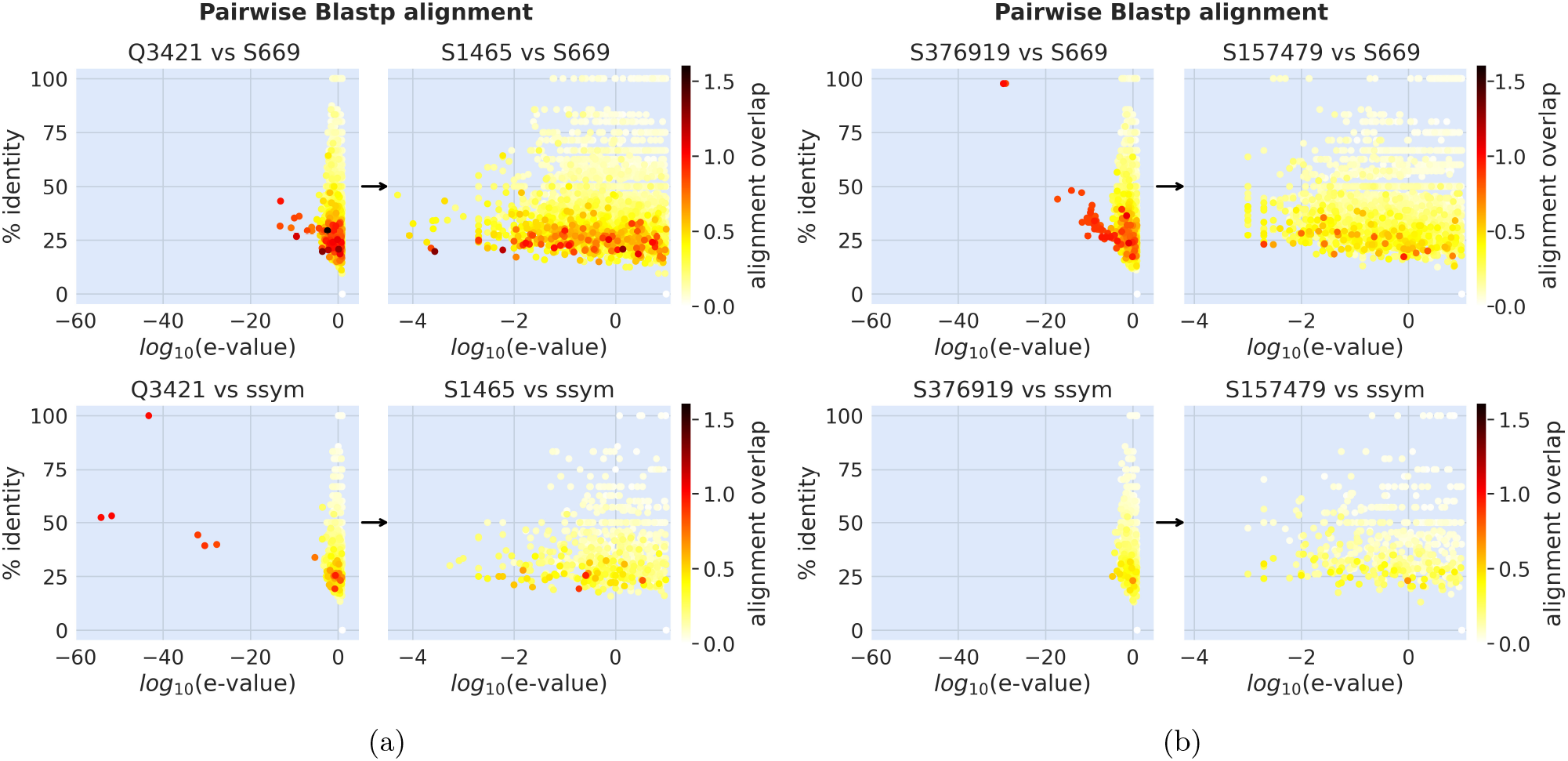
Pairwise BLASTp alignments (% identity, e-value and alignments overlap) of the training set against the *S669* (**top**) and *ssym* (**bottom**) test sets, before (**left**) and after (**right**) filtering by similarity. Each arrow indicates the transition from the original dataset to the filtered one. 1a: from *Q3421* to *S1465*. 1b: from *S376919* (the mega-scale dataset) to *S157479*.

Moreover, we utilize Multiple Sequence Alignments (MSAs) of protein families as a fundamental component of our analysis. Alignments are generated using HHblits [64] to search for homologous sequences within the Uniclust30 database [65]. To evaluate the train and test MSAs for any potential similarity among their sequences, we analyze pairs resulting in a BLASTp alignment overlap greater than 0.5. The MSA Hidden Markov Model (HMM) profiles of the 171 pairs, selected among all train/test datasets under consideration, are aligned using HHalign from the HHsuite3 package [66]. This analysis identifies four protein pairs (labeled by PDB-id) with anomalously low e-values (Figure 2a), prompting us to perform pairwise all-versus-all BLASTp alignments between sequences in their respective MSAs. Specifically, we focus on the MSAs of the two critical proteins from the large training set (1V1C and 2BTT) and we align their sequences against those in the MSA of the 2PRN protein of the *S669* test set (Figure 2b). Results reveal significant sequence similarity between certain pairs from the training and test MSAs. Similar findings are observed for the two critical proteins in the small training set (Figure 8 in Supporting Information). Based on these outcomes, we exclude these proteins from the training.

**Figure 2:**
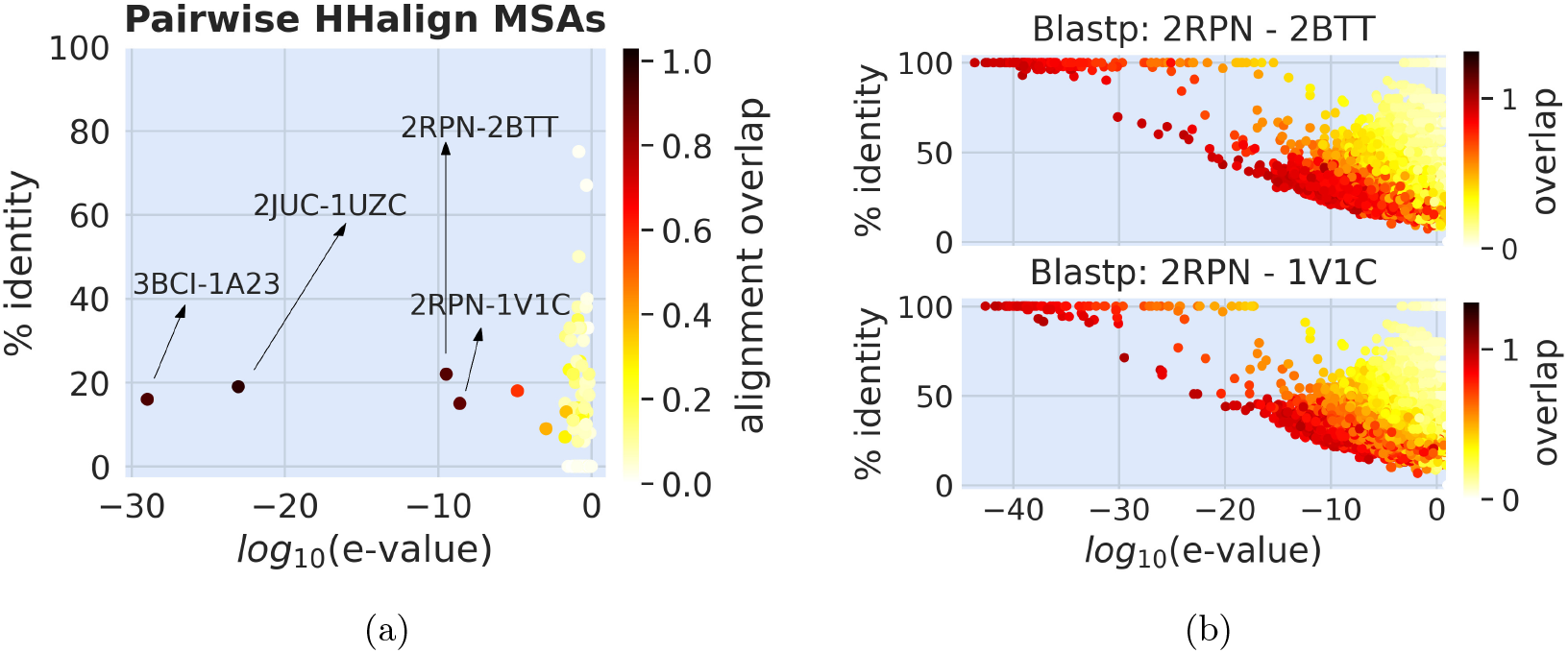
HHalign alignment of MSA HMM profiles between training and test sets (Fig. 2a). Pairwise Blastp alignment between sequences in the MSA of proteins 2BITT (Fig. 2b **top**) and 1V1C (Fig. 2b **bottom**) of the large training set, vs sequences in the MSA of 2PRN of the *S669* test set.

## 3 Methods

We present a framework for refining pre-trained PLMs through a supervised approach, harnessing their ability to encode biologically meaningful sequence representations. Our model is specifically designed to predict thermodynamic stability variations resulting from single amino acid substitutions in proteins. This is defined as the difference in the unfolding free energy between the mutated (M) and wild type (W) conformations, measured in kcal/mol:

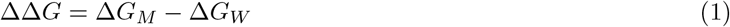

We conduct end-to-end fine-tuning of PLMs, with the PLM module independently processing mutated and wild-type sequences. Subsequently, a Multi-Layer Perceptron (MLP) takes in input the differences between mutated and wild-type representations extracted from the final hidden layer of the PLM. Specifically, we concatenate two vectors derived from the PLM’s last hidden layer: 1) the representation of the sequence, averaged across its length, and 2) the representation of the mutated amino acid position. The input to the MLP consists of the difference between these concatenated vectors for wild-type and mutated sequences. The model is trained to minimize the Mean Absolute Error between predicted and experimental ΔΔ*G* values. When training with the small dataset, we fine-tune three different protein language models: ESM-2 [58], ProstT5 [59], and MSA Transformer [57] (Figure 3), renaming them consequently. The initial pre-training weights are obtained from the HuggingFace Transformer Library [67]. The evaluation metrics are the Pearson’s correlation coefficient (r), the Mean Absolute Error (MAE), and the Root Mean Square Error (RMSE).

**Figure 3:**
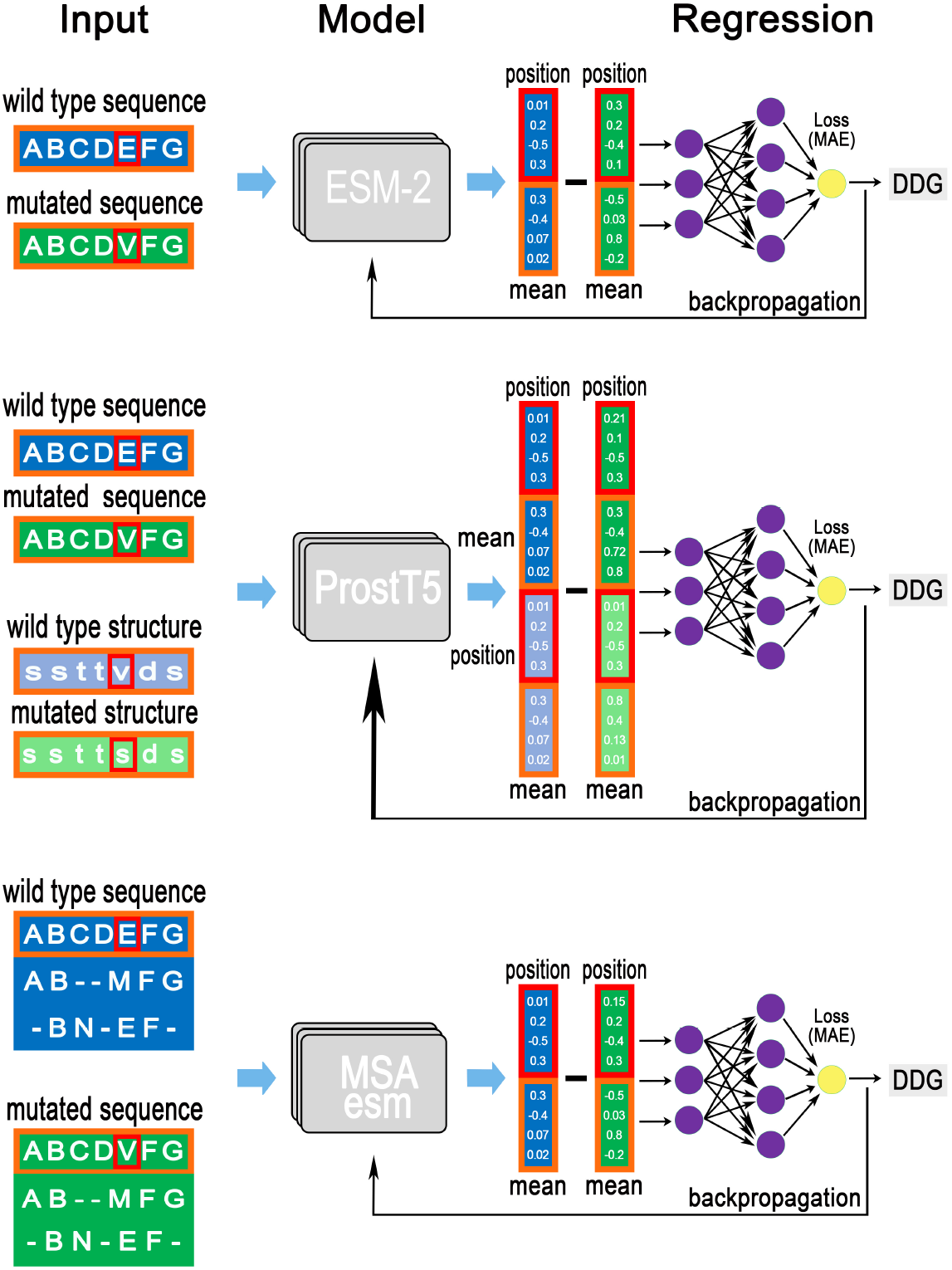
Models architectures: ESM2_ddG, ProstT5_ddG and MSAesm_ddG.

- **ESM2_ddG**: ESM-2 is one of the largest architectures among single sequence models and stands out for its role in structure prediction. We adopt the architecture with 640 million parameters and 36 layers. A comparable transfer learning approach involving the ESM-2 model is explored in a recent study [46], although it employs different training datasets.
- **ProstT5_ddG**: ProstT5, a bilingual protein language model featuring structure-aware embeddings, undergoes training on Foldseek’s 3Di structures [68] and amino acid sequences. Designed with an encoder-decoder transformer architecture, ProstT5 is tailored for translating between sequences and structures. In our methodology, we pre-process wild-type and mutated sequences by translating them into their corresponding 3Di structures. The ProstT5 encoder module is fine-tuned by processing both the sequences and the 3Di structures. We concatenate the resulting representations, feeding the difference between wild type and mutated ones into the regression head.
- **MSAesm_ddG**: The MSA Transformer introduces a structural modification compared to single sequence models, by processing a set of homologous aligned sequences. It achieves superior accuracy in contact prediction compared to individual sequence methods, while utilizing significantly fewer parameters. Notably, this model plays a pivotal role in the AlphaFold2 structure prediction algorithm [69]. In order to generate the mutated alignments, we substitute the mutated amino acid into the wild type. Only the query sequence representations are used to input the regression, similarly to the ESM-2_ddG architecture.

## 4 Results

### 4.1 Models and Datasets

We fine-tune ESM-2, ProstT5, and MSA Transformer models, trained with the small dataset (*S1413*), and evaluate performances on the *S669* test set (Table 1). This experimental setup serves as a reference for identifying the most effective protein language model among the assessed options. In the comparison, the MSA

**Table 1:** Performance of MSAesm_ddG, ESM2_ddG, and ProstT5_ddG, trained with the small *S1413* dataset.

| Models trained on *S1413* | *S669* test set |  |  |
| --- | --- | --- | --- |
|  | Pearson r | MAE | RMSE |
| MSAesm_ddG | 0.50 | 1.01 | 1.44 |
| ESM2_ddG | 0.47 | 1.03 | 1.45 |
| ProstT5_ddG | 0.41 | 1.08 | 1.52 |

Transformer emerges as the most promising model, while both the ESM2_ddG and the ProstT5_ddG methods lag behind it in terms of Pearson’s correlation coefficient. It is worth to mention that the optimized ESM_ddG demonstrates results on par with previous findings [46], if considering the more rigorous sequence similarity filtering applied to our training set. Ultimately, given the superior correlation outcome and its considerably lighter architecture compared to the other models under consideration, we decide to employ the fine-tuned MSA Transformer for subsequent analyses.

Furthermore, we leverage the smallest dataset to explore the impact on predictions of the ΔΔ*G* distribution symmetry within the training data. We invert the order of wild type and mutated sequences and change the sign of the ΔΔ*G* values to construct the reverse training set. For the symmetrical set, we include both the direct and reverse data. Training with these datasets, we note no significant impact on the measured accuracy (Table 4 in Supporting Information). This aligns with our expectations, as the inherent imbalance of the ΔΔ*G* distribution towards destabilizing mutations typically affects structure-based procedures that violate symmetry in their architectures. Relying on these observations, we choose to focus exclusively on direct mutations in the subsequent analyses. We analyze the behavior of various models by modifying the network configuration, to evaluate the importance of different components (Figure 4). We compare a baseline model without fine-tuning, bypassing the PLM training, and we conduct an ablation study using the sequence-mean and mutated position representation vectors separately as inputs for the MLP. Regarding fine-tuning, the MSA Transformer model shows an improvement, whereas the ESM2 model does not demonstrate similar gains. We also note that utilizing mutated position representations is the primary factor in enhancing performance, while incorporating the sequence-mean representation proves advantageous for both models. Furthermore, we fine-tune using the difference between the PLMs’ predicted probabilities of wild-type and mutated amino acids as MLP input. However, our findings indicate that this approach is insufficient to capture the complexity of the task, leading to a Pearson r of 0.33 and 0.35 for the MSA Transformer and ESM2 models, respectively.

**Figure 4:**
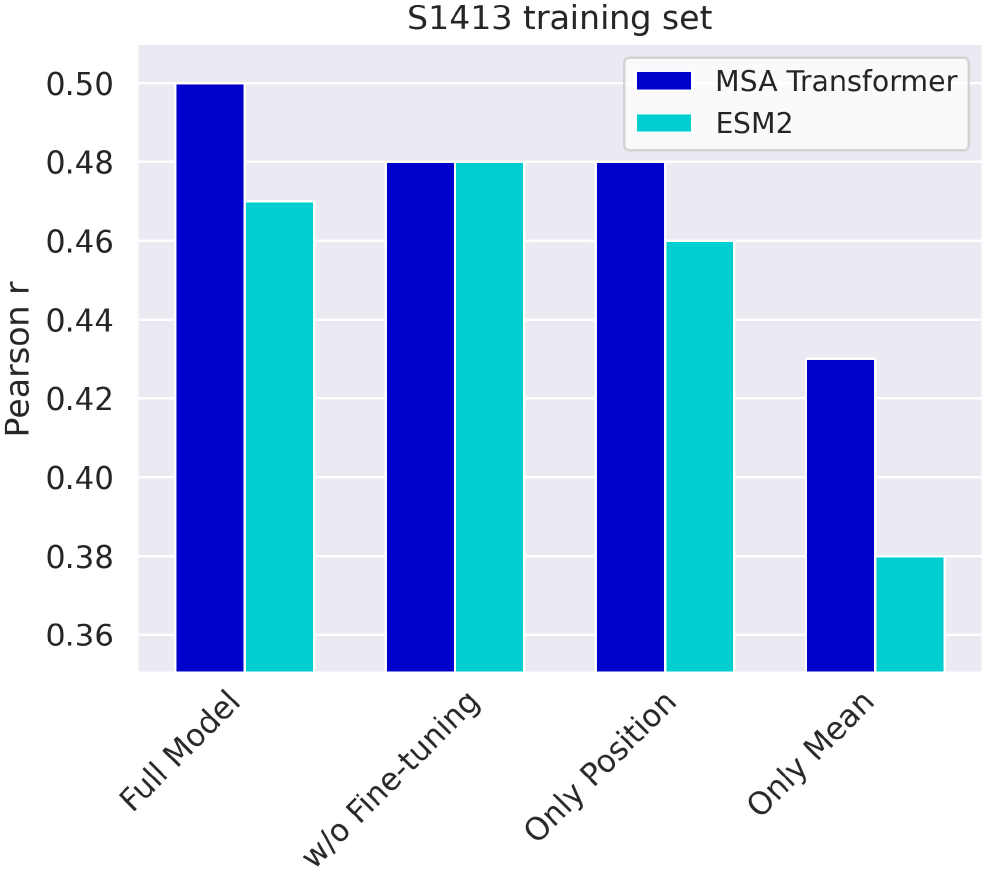
Pearson r for predictions on *S669* of optimized MSA Transformer and ESM2. Comparison of end-to-end trained models, baseline models (w/o fine-tuning), and ablation studies using only sequence-mean and only position representations as MLP input.

We trained the MSAesm_ddG model with both the large dataset (*S155329*) and the much smaller, designed dataset (*S3648*). Our findings demonstrate the appreciable advantage of training with the mega-scale data compared to the previously adopted *S1413* (Figure 5). Interestingly, despite its considerably smaller size, training with the designed dataset yields comparable accuracy, resulting in the lowest MAE values. This can be due to its extended range and variety of proteins, with respect to the small and large datasets.

**Figure 5:**
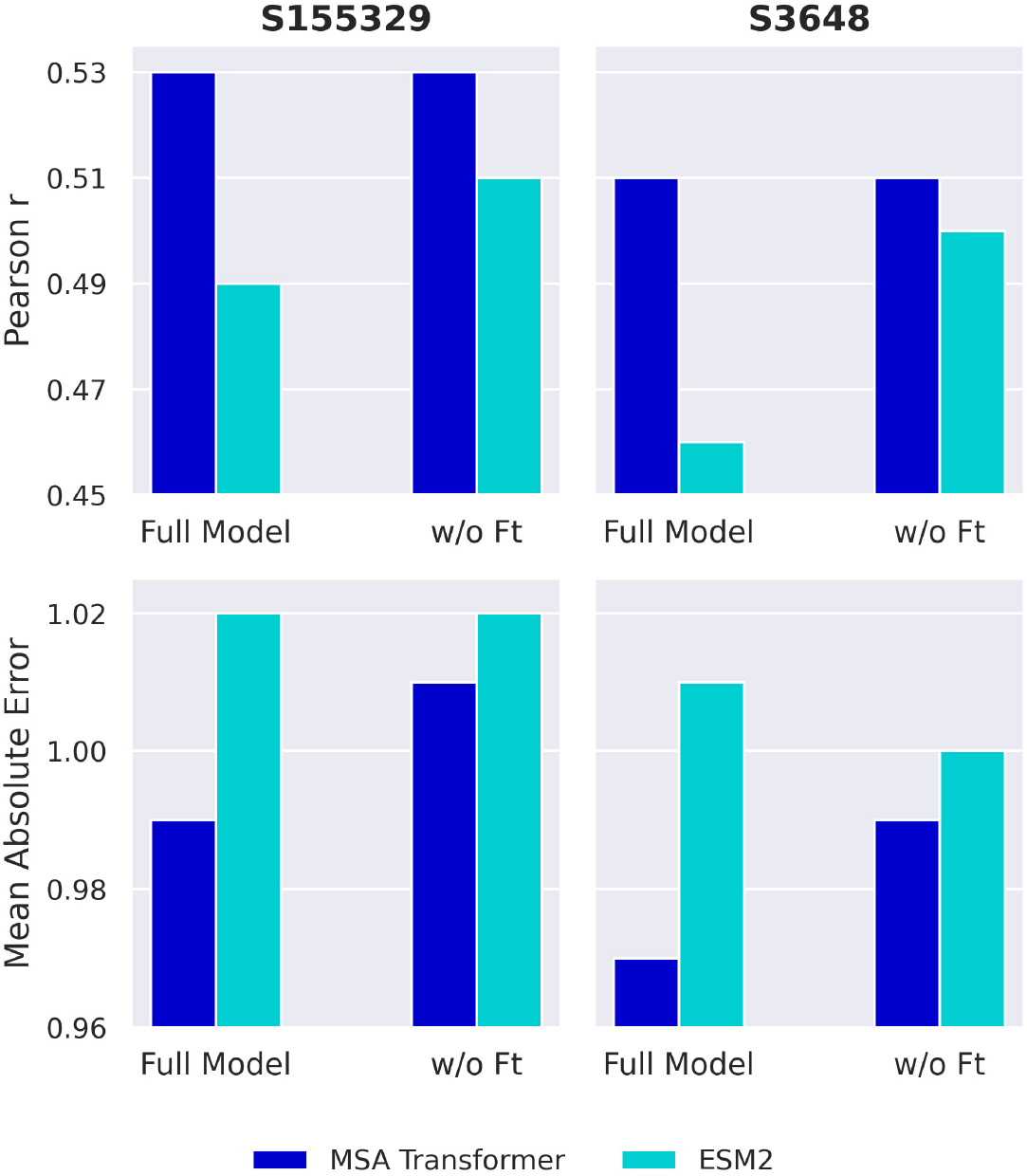
Pearson r (**top**) and Mean Absolute Error (**bottom**) for predictions on *S669* for the fine-tuned and baseline (w/o ft) MSA Transformer and ESM2 models, trained with datasets *S155329* (**left**) and *S3648* (**right**).

For subsequent comparisons with established methods we prioritize the MSAesm_ddG model, trained on the large dataset. The motivation for this choice is twofold: 1) MSAesm_ddG shows superior efficacy with respect to the other models under investigation and 2) the large training dataset offers the opportunity to evaluate the performance on the *ssym* data, not used during validation. It should be emphasized that, while the end-to-end procedure performs very well, training solely the MLP block can still produce remarkably good results. This underscores the effectiveness of pre-training in imparting the evolutionary knowledge required to adeptly address the task. Notably, fine-tuning enhances the performance of the MSA transformer, as evidenced by the reduction in MAE, but it proves to be detrimental for ESM2, highlighting the variability in its effectiveness across different models. These observations necessitate additional experiments to both identify and potentially explain the transformation in representations resulting from the fine-tuning process.

### 4.2 Comparison of prediction methods

We evaluate the efficacy of the MSAesm_ddG model, trained extensively on the mega-scale dataset, through a comprehensive comparison with various models as outlined in [45]. The analysis is performed using the *S669* test set, specifically designed to prevent overfitting for all methods (Table 2). Performance metrics for recently released algorithms are sourced from their respective manuscripts [40,46,51]. The techniques under comparison encompass a variety of approaches, including classical and machine-learning (ML) based methods. Only a few methods focus solely on sequence information, while the majority incorporate structural data.

**Table 2:** Methods performance on the *S669* test set.

|  | Method | *S669* test set |  |  | Method Description | Dataset |
| --- | --- | --- | --- | --- | --- | --- |
|  |  | r | MAE | RMSE |  |  |
| Machine Learning<br>sequence-based | MSAesm_ddG | **0.53** | **0.99** | **1.41** | MSA Transformer + MLP | S155329<sup>†</sup> |
|  | PROSTATA | 0.49 | 1.00 | 1.45 | ESM2 + MLP | S5251<sup>†</sup> |
|  | INPS-Seq | 0.43 | 1.09 | 1.52 | Support Vector Regression | S2648* |
|  | ACDC-NN-Seq | 0.42 | 1.08 | 1.53 | Convolutional Differential NN | S2648*, Varibench |
| sequence-based | DDGun | 0.41 | 1.25 | 1.72 | Linear combination of evo-scores | S2648*, Varibench |
| Machine Learning<br>structure-based | ThermoMPNN | 0.43 | - | 1.52 | Graph NN+Attention block+MLP | S216920<sup>†</sup> |
| | RaSP | 0.39 | 1.14 | 1.63 | 3D Convolutional + FFNN | S257695 (Rosetta $\Delta\Delta G$) |
|  | Dynamut2 | 0.34 | 1.15 | 1.58 | Random Forest Regression | S2648* |
|  | ACDC-NN | 0.46 | 1.05 | 1.49 | Convolutional Differential NN | S2648*, Varibench |
|  | ThermoNet | 0.39 | 1.17 | 1.62 | 3D Convolutional NN | Q3488* |
|  | PremPS | 0.41 | 1.08 | 1.50 | Random Forest Regression | S2648* |
|  | Dynamut | 0.41 | 1.19 | 1.60 | Random Forest Regression | S2648* |
|  | INPS3D | 0.43 | 1.07 | 1.50 | Support Vector Regression | S2648* |
|  | DUET | 0.41 | 1.10 | 1.52 | Support Vector Regression | S2297* |
|  | MAESTRO | 0.50 | 1.06 | 1.44 | Multi-agent of models ensemble | S2298*, S1925**, S1765* |
|  | PoPMuSiC | 0.41 | 1.09 | 1.51 | LR of statistical potentials | S2648* |
|  | I-Mutant3.0 | 0.36 | 1.12 | 1.52 | Support Vector Machine | S3246** |
| structure-based | DDGun3D | 0.43 | 1.11 | 1.60 | Linear combination of local features | S2648*, Varibench |
|  | SDM | 0.41 | 1.26 | 1.67 | Environment-specific substitutions | S388* |
|  | Rosetta | 0.39 | 2.08 | 2.70 | Energy functions + sampling | PDB |
|  | FoldX | 0.22 | 1.56 | 2.30 | Empirical Force Field | S2648*, Varibench |

<sup>†</sup>Data extracted from the megascale database [60].
\*Data extracted from ProTherm version 4.0 [70]. \*\*Data extracted from ProTherm [71].

- **Machine Learning methods**. *Sequence-based* : ACDC-NN-Seq [41], INPS-Seq [42]. *Structure-based* : I-Mutant3.0 [29], PoPMuSiC [30], MAESTRO DUET [31], [32], INPS3D [33], Dynamut [34], PremPS [35], ThermoNet [36], ACDC-NN [37], Dynamut2 [38], RaSP [40]. **PLM-based methods**. *Sequence-based* : PROSTATA [46]. *Structure-based* : ThermoMPNN [51].
- **Classical methods**. *Sequence-based* : DDGun [28]. *Structure-based* : FoldX [25], Rosetta [26], SDM [27], DDGun3D [28].

Additionally, we provide results on the *ssym* test set (Table 3), where we compare the performance of MSAesm_ddG with that of another PLM-based model [46], given the authors’ assertion of no data leakage for the *ssym* test set.

**Table 3:** Performance of two PLM-based methods on the *ssym* test set.

| Method | *ssym* test set |  |  |
| --- | --- | --- | --- |
|  | Pearson r | MAE | RMSE |
| MSAesm_ddG | **0.59** | **0.90** | **1.29** |
| PROSTATA | 0.51 | - | 1.40 |

The results underscore the robustness and effectiveness of the developed method, establishing it as a leading approach for predicting the impact of mutations on protein stability. Furthermore, MSAesm_ddG outperforms a similar PLM-based approach that involves fine-tuning the single-sequence ESM-2 model, emphasizing the crucial role of integrating evolutionary information into the model architecture.

### 4.3 Impact of protein fold class on predictions

We present illustrative results for proteins with more than 20 mutations in both the *S669* (Figure 6a) and *ssym* (Figure 6b) datasets. The majority of predictions for these proteins exhibit a Pearson correlation coefficient above 0.6, while only 30% show a lower correlation with the experimental ΔΔ*G* values. Remarkably, the protein with the lowest correlation (2JIE) comprises a large number of residues, forcing to restrict the inference to only a limited number of homologous due to memory usage constraints. This limitation could potentially contribute to the poorer performance of MSAesm_ddG for this specific protein. Overall, these findings highlight how the model’s accuracy is distinctly influenced by the specific protein under consideration, prompting further investigation in this regard.

**Figure 6:**
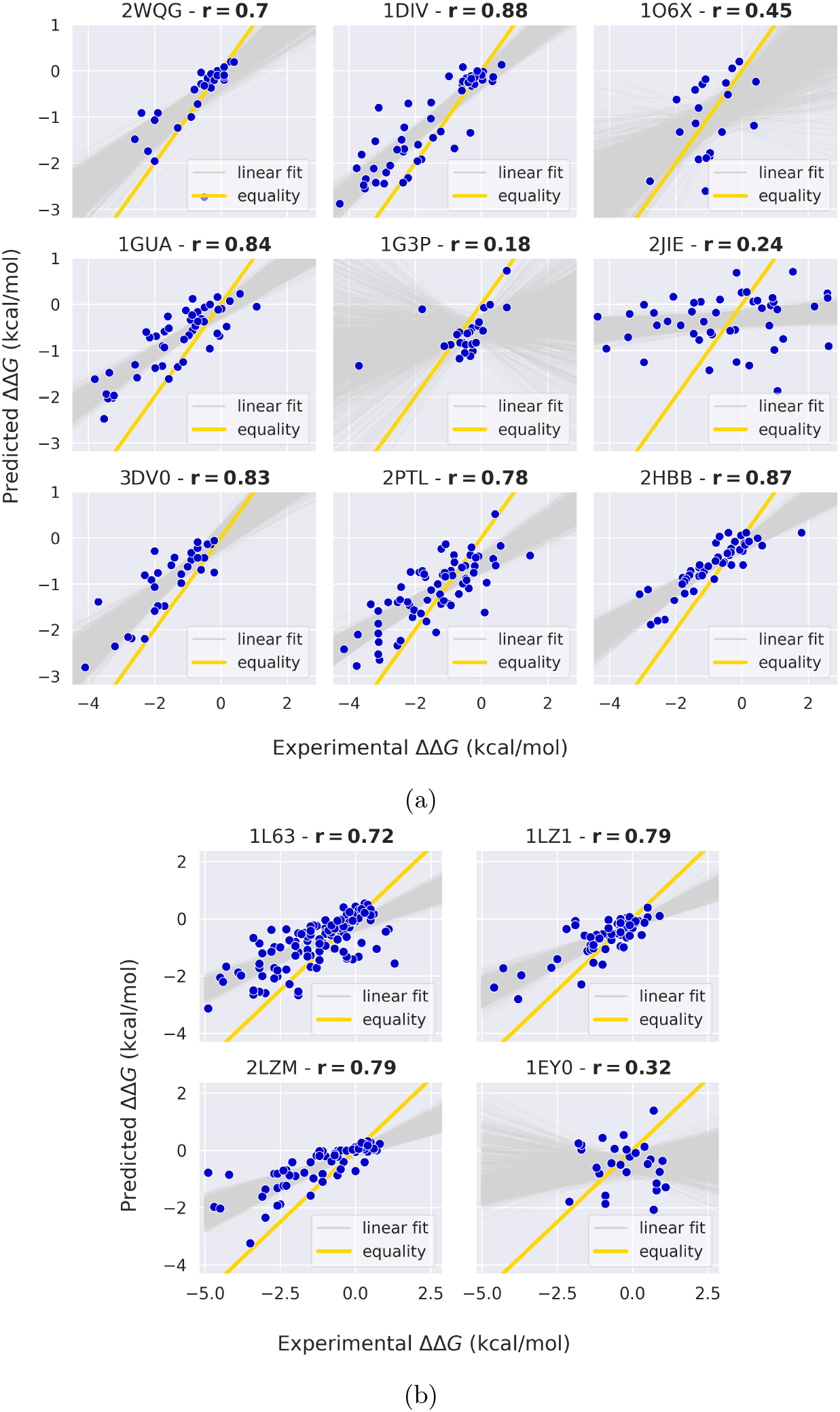
Predicted and experimental ΔΔ*G* of proteins presenting more than 20 mutations. 6a: *S669* test set. 6b: *ssym* test set. The linear fit over 1000 bootstrapped samples in *gray* shows the trend of the shift between predictions and experiments.

Examining the effective slope of the linear fit between predictions and experiments, we measure a consistent shift between the two across most depicted proteins. This effect arises from the diverse nature of the training and test ΔΔ*G* distributions, stemming from different experimental conditions. In particular, the training distribution exhibits less prominent tails compared to the test data (Figure 7), which reflects into a narrower range of predicted values with respect to the experimental ΔΔ*G* in the test set. By excluding the tails from the experimental *S669* distribution and retaining the range of ΔΔ*G* values from -4 to 2 kcal/mol, we obtain a slightly reduced test set (*S625*). This reduction results in significantly improved performance metrics (r=0.54, MAE=0.82, RMSE=1.08), demonstrating the sensitivity of our model to the distinct ranges of ΔΔ*G* values spanned by the training and test distributions.

**Figure 7:**
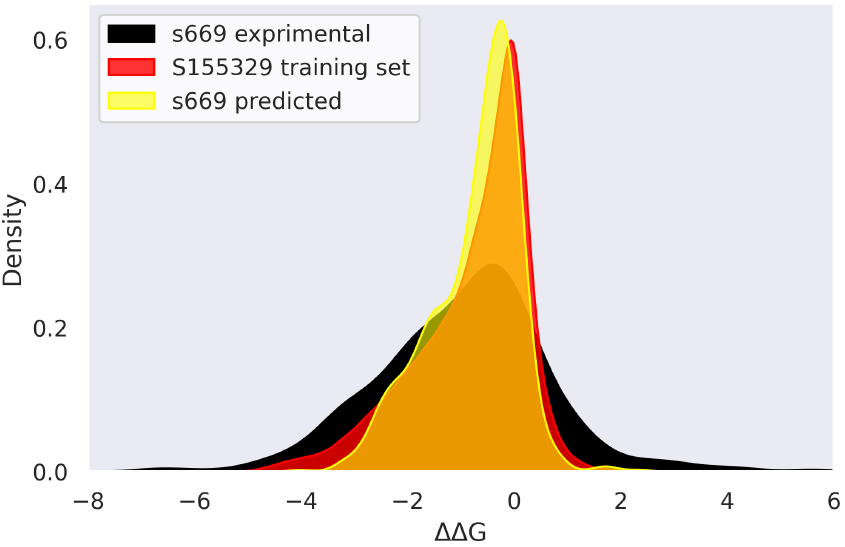
ΔΔ*G* probability density of the experimental S669, the predicted S669 and the large training set.

To delve deeper into the factors influencing predictive performance, we focus on proteins with a minimum of 5 mutations in the test sets and categorize them through the CATH fold class annotation [72]. Predictions are classified as positive cases if they demonstrate a correlation above the Pearson correlation coefficient measured on the full test set (21 cases), and negative if they fall below it (15 cases). Results reveal that among the analyzed cases, positive examples consist of 9.5% mainly beta, 38% mainly alpha, and 52.5% alpha/beta secondary structure motifs. Conversely, negative proteins comprise 47% mainly beta, 27% mainly alpha, and 27% alpha/beta secondary structure motifs. Of particular note is the observation that the number of superfamilies annotated as mainly alpha structures in CATH is nearly twice as large as those annotated as mainly beta. From this observation, we hypothesize that the prevalence of mainly alpha secondary structures among protein families could contribute to higher uncertainty in the model’s predictions for mainly beta annotated proteins, which are less represented in the pre-training of the MSA Transformer.

## 5 Discussion

The advent of Protein Language Models as robust instruments for deciphering the language embedded within amino acid sequences has profoundly influenced the landscape of structural biology. In this study, we delved into the efficacy of PLMs to forecast thermodynamic stability changes induced by single amino acid mutations, introducing an innovative approach that leverages the MSA Transformer model. In order to actively address the pervasive issue of overfitting, we filter training sequences adopting a stringent approach based on their resemblance to test sequences. First of all, our findings reveal the superior performance of the optimized MSA Transformer model compared to other PLMs such as ESM-2 and ProstT5. Through comprehensive comparisons of frameworks and training strategies, we find that fine-tuning benefits the MSA Transformer model, with ablation studies validating the robustness of the designed architecture. While training with a mega-scale dataset yields optimal results, we observe comparable accuracy when employing a smaller dataset designed to encompass a broader range of proteins. This suggests that the injection of diversity into the training dataset, even with a smaller mutation count, significantly amplifies the model’s generalization capabilities. Nonetheless, our MSA-based model consistently outperforms classical and machine learning-based methods that span both sequence-based and structure-based approaches. Through protein-wise analysis, we unveil intriguing trends in prediction accuracy across diverse protein structures. Notably, we observe a discernible dependency of prediction accuracy on the specific protein under consideration, suggesting that structural peculiarities inherent to individual proteins play a crucial role in shaping predictive outcomes. Positive instances, characterized by a higher correlation between predicted and experimental stability changes, are primarily associated with alpha/beta secondary structure motifs, while negative instances, exhibiting lower correlation, are predominantly linked with primarily beta structures. This underscores the importance of considering protein-specific features in stability prediction models and warrants further investigation into the nuanced relationship between protein structure and prediction accuracy. Another pivotal aspect of our study lies in the comparison with established prediction methods. A key advantage of the MSA Transformer lies in its proficiency at leveraging co-evolution signals inherent in homologous sequences within Multiple Sequence Alignments. These signals, reflecting conserved residue interactions throughout evolutionary history, provide valuable insights into the functional and structural constraints governing proteins. By integrating this rich source of evolutionary information, the MSA Transformer effectively discriminates between stabilizing and destabilizing mutations, even in cases where conventional methods struggle. Moreover, the model benefits from its robust architecture, which is specifically designed to handle large-scale sequence data and capture long-range dependencies. The self-attention mechanism employed in the transformer architecture allows the model to focus on relevant regions of the sequence and determine the importance of different amino acids based on their contextual information within the alignment. This attention mechanism enables to effectively discern changes in sequence patterns that are indicative of stability-altering mutations. Unlike traditional machine learning approaches, reliant on domain-specific features or handcrafted representations, our model learns directly from raw sequence data, rendering it highly adaptable to diverse protein structures and functional contexts. This adaptability empowers the model to outperform existing methods, albeit contingent upon the availability of homologous sequences. In addition to sequence coverage, it is crucial to emphasize that the specific techniques and databases used for constructing Multiple Sequence Alignments can significantly influence results. In this study, we adopted the same strategy as the MSA Transformer model’s pre-training, searching for homologs in a highly non-redundant database to enhance the co-evolutionary signal in our alignments, potentially aiding the learning process. In summary, our study advances the field by demonstrating the effectiveness of fine-tuning protein language models using evolutionary information for predicting stability changes induced by mutations. Looking ahead, further advancements in PLM-based architectures, along with the increasing availability of comprehensive sequence databases, are set to drive continued progress in this domain, facilitating the design of more stable and functional proteins for diverse biomedical and biotechnological applications.

## 7 Acknowledgments and Funding

The authors acknowledge the AREA Science Park supercomputing platform ORFEO made available for conducting the research reported in this paper and the technical support of the Laboratory of Data Engineering staff. F.C., M.C., A.A., and A.C. were supported by the European Union – NextGenerationEU within the project PNRR ”PRP@CERIC” IR0000028 - Mission 4 Component 2 Investment 3.1 Action 3.1.1.

## Supporting Information

**Figure 8:**
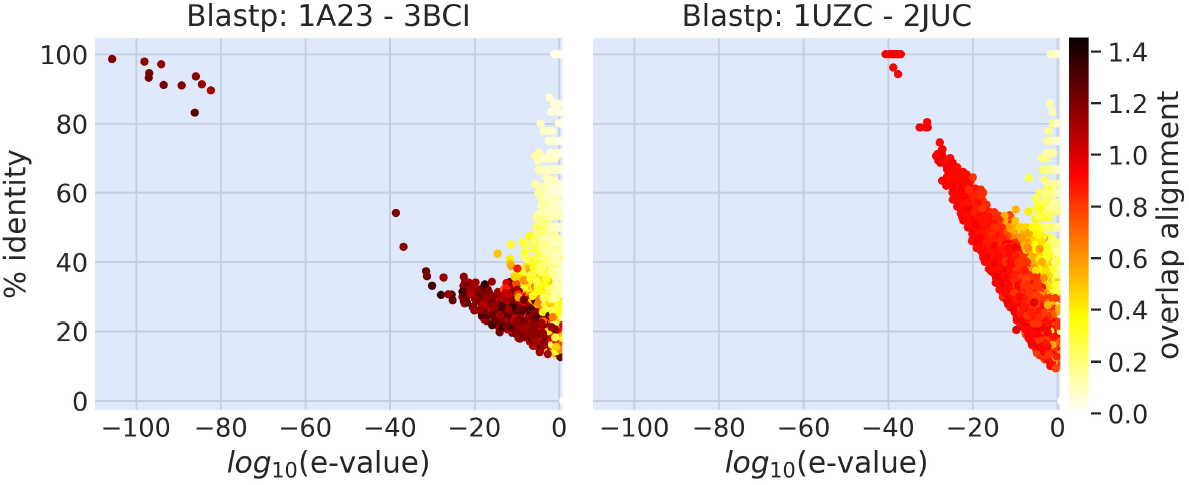
Pairwise Blastp alignment between sequences in MSA of 1A23 (*S1413* training set) vs 3BCI (small test set) (**left**) and 1UZC (*S1413* training set) vs 2JUC (small test set) (**right**).

**Figure 9:**
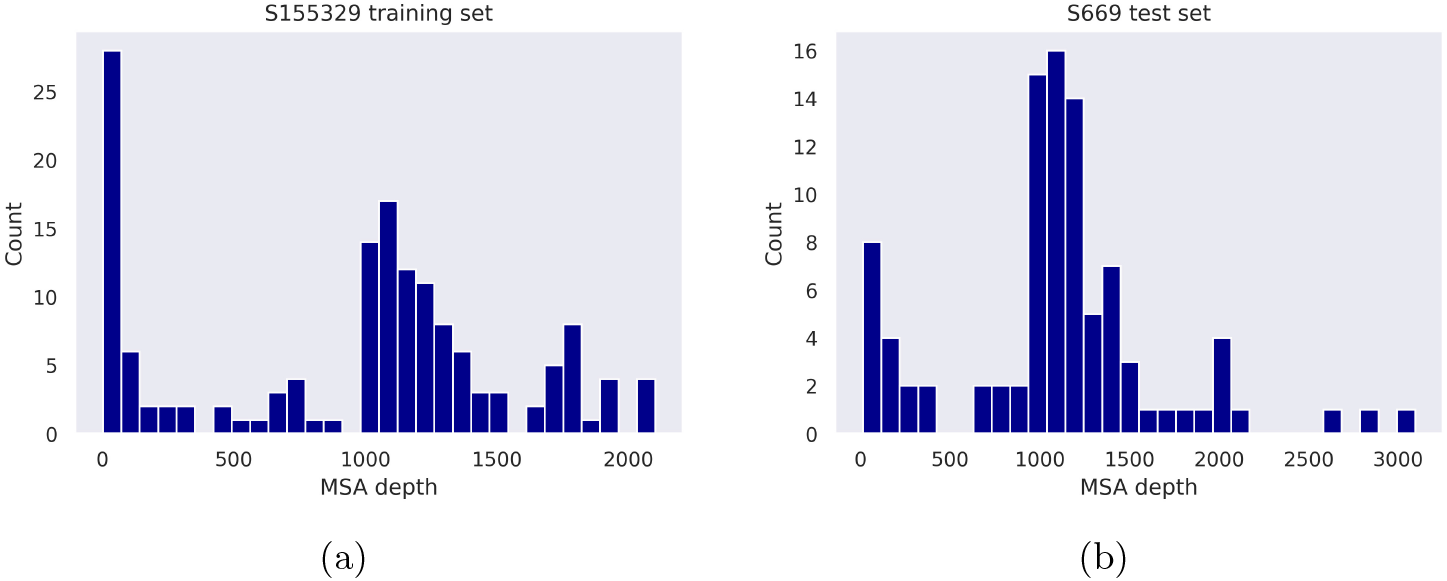
MSA depth (number of sequences) in the *S155329* training set (9a) and in the *S669* test set (9b).

**Figure 10:**
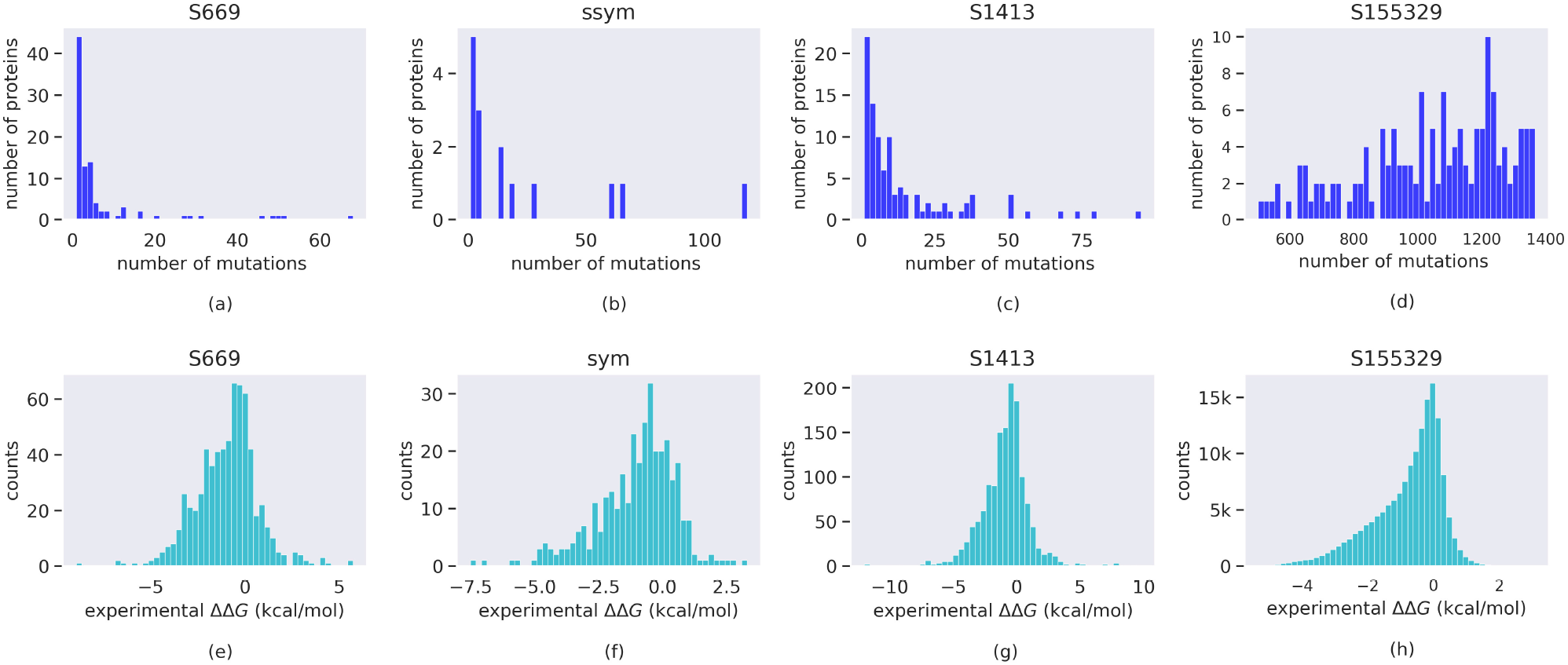
Number of proteins vs number of mutations (a-d) and ΔΔ*G* distribution (e-h) for all datasets.

### Tuning Hyperparameters

The Multi-Layer Perceptron block consists of one hidden layer with size 768, followed by a ReLU activation and a dropout layer (with probability *p* of an element to be zeroed set to *p* = 0.2) that feeds into the final linear layer. We adopt a PyTorch implementation with AdamW optimizer (`torch.optim.AdamW` with default parameters, apart from the learning rate) and OneCycleR learning-rate scheduler(`torch.optim.lr_scheduler.OneCycleLR`). The learning rate is selected for each model and training dataset by means of a grid search with lr ∈ 0.5×10^−5^, 10^−5^, 0.5×10^−4^, 10^−4^, 0.5×10^−3^, 10^−3^, 0.5×10^−2^, 10^−2^. In order to reduce the memory footprint, the PyTorch Automatic Mixed Precision is enabled by means of `torch.autocast` (specifically using `dtype=torch.float16`) in combination with the gradient scaling function `torch.cuda.amp.GradScaler()`. The gradient norm is clipped using `torch.nn.utils.clip grad norm` with `max norm` = 10, to prevent possible exploding gradient problems.

We set a maximum number of 20 epochs for the training in combination with an early stopping criterion. The latter consists in saving the model parameters associated to the epoch with lowest MAE on the validation set, and stopping the training if the MAE on the validation is not decreasing for the next 5 epochs. For training with both the small and designed datasets, we validate on the *ssym* set, which does not overlap with both the training and the test sequences (Figure 11a). When training with the large dataset, a customized validation set is constructed (*S1030*) by excluding from the small dataset sequences that are similar to those in the large dataset (Figure 11b), following the criterion stated in the main text.

Using the PyTorch DistributedDataParallel class (`torch.nn.parallel.DistributedDataParallel`), the fine-tuning is performed on 16 nodes of the LEONARDO Booster partition at CINECA National Supercomputing Center, each consisting of a single socket 32 cores Intel Xeon Platinum 8358 (2.60GHz), together with×4 NVIDIA A100 GPUs (64GB HBM2e and NVLink 3.0) and 512 GB DDR4 (3200 MHz) RAM. The internal network is provided by NVIDIA Mellanox Infiniband HDR DragonFly+ 200 Gbps.BB We consider a batch size = 1 for a single GPU, so the overall effective batch size _eff_ = 64. Finally, we load the model parameters for the inference on the test sets on a single GPU.

**Figure 11:**
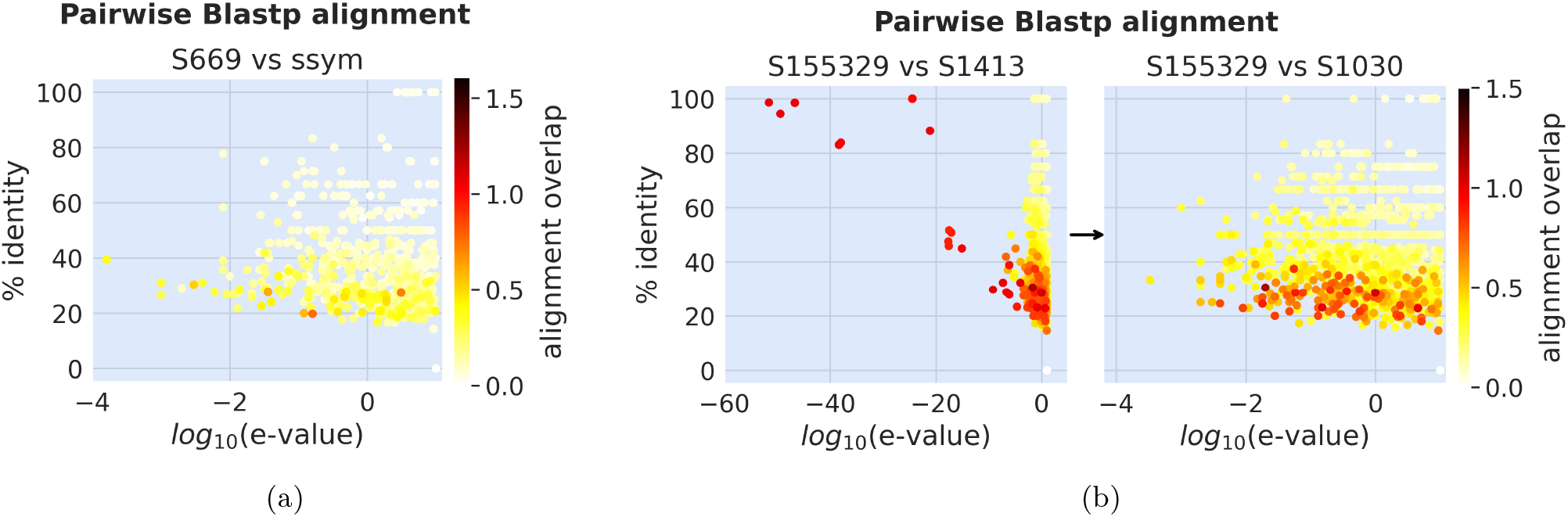
Pairwise BLASTp alignments (% identity, e-value and alignments overlap). 11a: *S669* test set against the ssym set. 11b: *S155329* training set against the *S1413* dataset, before (**left**) and after (**right**) filtering by similarity to obtain the *S1030* dataset (used for validation in the *S155329* training).

**Table 4:**
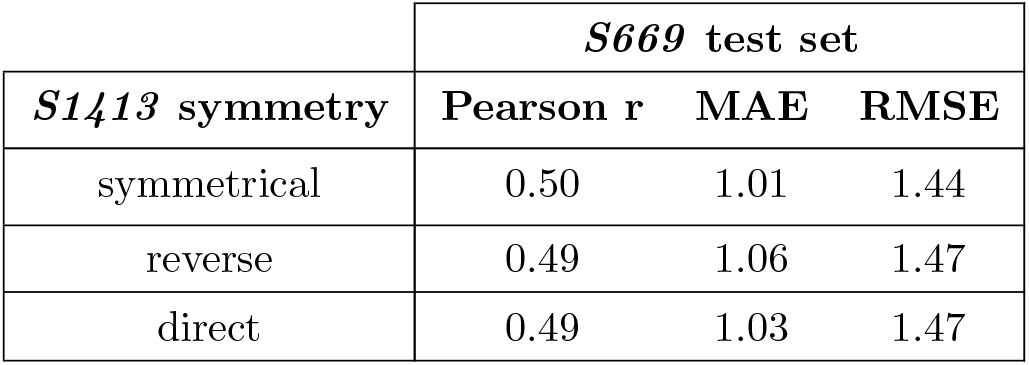
MSAesm_ddG model trained with the symmetrical, reverse and direct ΔΔ*G* distributed *S1413*.

**Table 5:** MSAesm_ddG fine-tuned and baseline, trained with different datasets.

| Training sets | *S669* test set - Ft |  |  | *S669* test set - w/o Ft |  |  |
| --- | --- | --- | --- | --- | --- | --- |
|  | Pearson r | MAE | RMSE | Pearson r | MAE | RMSE |
| *S155329* | 0.53 | 0.99 | 1.41 | 0.53 | 1.00 | 1.43 |
| *S3648* | 0.51 | 0.97 | 1.41 | 0.51 | 0.99 | 1.41 |
| *S1413* | 0.50 | 1.01 | 1.44 | 0.48 | 1.03 | 1.44 |

**Table 6:** ESM2_ddG fine-tuned and baseline, trained with different datasets.

|  | *S669* test set - Ft |  |  | *S669* test set - w/o Ft |  |  |
| --- | --- | --- | --- | --- | --- | --- |
| Training sets | Pearson r | MAE | RMSE | Pearson r | MAE | RMSE |
| *S155329* | 0.49 | 1.02 | 1.47 | 0.51 | 1.02 | 1.46 |
| *S3648* | 0.46 | 1.01 | 1.46 | 0.50 | 1.00 | 1.43 |
| *S1413* | 0.48 | 1.03 | 1.45 | 0.47 | 1.03 | 1.45 |

## References

[1] Ashish Vaswani, Noam Shazeer, Niki Parmar, Jakob Uszkoreit, Llion Jones, Aidan N Gomez, Lukasz Kaiser, and Illia Polosukhin. Attention is all you need. Advances in neural information processing systems, 30, 2017.

[2] Jesse Vig, Ali Madani, Lav R Varshney, Caiming Xiong, Richard Socher, and Nazneen Fatema Rajani. Bertology meets biology: interpreting attention in protein language models. arXiv preprint 2006.15222, 2020.

[3] Tristan Bepler and Bonnie Berger. Learning the protein language: Evolution, structure, and function. Cell systems, 12(6):654–669, 2021.

[4] Nicki Skafte Detlefsen, Søren Hauberg, and Wouter Boomsma. Learning meaningful representations of protein sequences. Nature communications, 13(1):1914, 2022.

[5] Lucrezia Valeriani, Diego Doimo, Francesca Cuturello, Alessandro Laio, Alessio Ansuini, and Alberto Cazzaniga. The geometry of hidden representations of large transformer models. Advances in Neural Information Processing Systems, 36, 2024.

[6] Michael Heinzinger, Ahmed Elnaggar, Yu Wang, Christian Dallago, Dmitrii Nechaev, Florian Matthes, and Burkhard Rost. Modeling aspects of the language of life through transfer-learning protein sequences. BMC bioinformatics, 20(1):1–17, 2019.

[7] Roshan Rao, Joshua Meier, Tom Sercu, Sergey Ovchinnikov, and Alexander Rives. Transformer protein language models are unsupervised structure learners. Biorxiv, pages 2020–12, 2020.

[8] Alexander Rives, Joshua Meier, Tom Sercu, Siddharth Goyal, Zeming Lin, Jason Liu, Demi Guo, Myle Ott, C Lawrence Zitnick, Jerry Ma, et al. Biological structure and function emerge from scaling unsupervised learning to 250 million protein sequences. Proceedings of the National Academy of Sciences, 118(15):e2016239118, 2021.

[9] Brian Hie, Ellen D Zhong, Bonnie Berger, and Bryan Bryson. Learning the language of viral evolution and escape. Science, 371(6526):284–288, 2021.

[10] Joshua Meier, Roshan Rao, Robert Verkuil, Jason Liu, Tom Sercu, and Alex Rives. Language models enable zero-shot prediction of the effects of mutations on protein function. Advances in Neural Information Processing Systems, 34:29287–29303, 2021.

[11] Brian L Hie, Kevin K Yang, and Peter S Kim. Evolutionary velocity with protein language models predicts evolutionary dynamics of diverse proteins. Cell Systems, 13(4):274–285, 2022.

[12] Nicole N Thadani, Sarah Gurev, Pascal Notin, Noor Youssef, Nathan J Rollins, Daniel Ritter, Chris Sander, Yarin Gal, and Debora S Marks. Learning from prepandemic data to forecast viral escape. Nature, pages 1–8, 2023.

[13] Nobuhiko Tokuriki, Francois Stricher, Luis Serrano, and Dan S Tawfik. How protein stability and new functions trade off. PLoS computational biology, 4(2):e1000002, 2008.

[14] Nobuhiko Tokuriki and Dan S Tawfik. Protein dynamism and evolvability. Science, 324(5924):203–207, 2009.

[15] Jesse D Bloom, Sy T Labthavikul, Christopher R Otey, and Frances H Arnold. Protein stability promotes evolvability. Proceedings of the National Academy of Sciences, 103(15):5869–5874, 2006.

[16] Konstantin B Zeldovich, Peiqiu Chen, and Eugene I Shakhnovich. Protein stability imposes limits on organism complexity and speed of molecular evolution. Proceedings of the National Academy of Sciences, 104(41):16152–16157, 2007.

[17] James W Bryson, Stephen F Betz, Helen S Lu, Daniel J Suich, Hongxing X Zhou, Karyn T O’Neil, and William F DeGrado. Protein design: a hierarchic approach. Science, 270(5238):935–941, 1995.

[18] Michele Vendruscolo, Amos Maritan, and Jayanth R Banavar. Stability threshold as a selection principle for protein design. Physical review letters, 78(20):3967, 1997.

[19] Gabriel J Rocklin, Tamuka M Chidyausiku, Inna Goreshnik, Alex Ford, Scott Houliston, Alexander Lemak, Lauren Carter, Rashmi Ravichandran, Vikram K Mulligan, Aaron Chevalier, et al. Global analysis of protein folding using massively parallel design, synthesis, and testing. Science, 357(6347):168–175, 2017.

[20] Zhen Wang and John Moult. Snps, protein structure, and disease. Human mutation, 17(4):263–270, 2001.

[21] Pauline C Ng and Steven Henikoff. Predicting deleterious amino acid substitutions. Genome research, 11(5):863–874, 2001.

[22] Peng Yue, Zhaolong Li, and John Moult. Loss of protein structure stability as a major causative factor in monogenic disease. Journal of molecular biology, 353(2):459–473, 2005.

[23] AndreasC Joerger and Alan R Fersht. Structure–function–rescue: the diverse nature of common p53 cancer mutants. Oncogene, 26(15):2226–2242, 2007.

[24] Boris Reva, Yevgeniy Antipin, and Chris Sander. Predicting the functional impact of protein mutations: application to cancer genomics. Nucleic acids research, 39(17):e118–e118, 2011.

[25] Joost Schymkowitz, Jesper Borg, Francois Stricher, Robby Nys, Frederic Rousseau, and Luis Serrano. The foldx web server: an online force field. Nucleic acids research, 33(suppl 2):W382–W388, 2005.

[26] Rebecca F Alford, Andrew Leaver-Fay, Jeliazko R Jeliazkov, Matthew J O’Meara, Frank P DiMaio, Hahnbeom Park, Maxim V Shapovalov, P Douglas Renfrew, Vikram K Mulligan, Kalli Kappel, et al. The rosetta all-atom energy function for macromolecular modeling and design. Journal of chemical theory and computation, 13(6):3031–3048, 2017.

[27] Arun Prasad Pandurangan, Bernardo Ochoa-Montano, David B Ascher, and Tom L Blundell. Sdm: a server for predicting effects of mutations on protein stability. Nucleic acids research, 45(W1):W229–W235, 2017.

[28] Ludovica Montanucci, Emidio Capriotti, Giovanni Birolo, Silvia Benevenuta, Corrado Pancotti, Dennis Lal, and Piero Fariselli. Ddgun: an untrained predictor of protein stability changes upon amino acid variants. Nucleic Acids Research, 50(W1):W222–W227, 2022.

[29] Emidio Capriotti, Piero Fariselli, and Rita Casadio. I-mutant2. 0: predicting stability changes upon mutation from the protein sequence or structure. Nucleic acids research, 33(suppl 2):W306–W310, 2005.

[30] Yves Dehouck, Jean Marc Kwasigroch, Dimitri Gilis, and Marianne Rooman. Popmusic 2.1: a web server for the estimation of protein stability changes upon mutation and sequence optimality. BMC bioinformatics, 12(1):1–12, 2011.

[31] Douglas EV Pires, David B Ascher, and Tom L Blundell. Duet: a server for predicting effects of mutations on protein stability using an integrated computational approach. Nucleic acids research, 42(W1):W314–W319, 2014.

[32] Josef Laimer, Heidi Hofer, Marko Fritz, Stefan Wegenkittl, and Peter Lackner. Maestro-multi agent stability prediction upon point mutations. BMC bioinformatics, 16(1):1–13, 2015.

[33] Castrense Savojardo, Piero Fariselli, Pier Luigi Martelli, and Rita Casadio. Inps-md: a web server to predict stability of protein variants from sequence and structure. Bioinformatics, 32(16):2542–2544, 2016.

[34] Carlos HM Rodrigues, Douglas EV Pires, and David B Ascher. Dynamut: predicting the impact of mutations on protein conformation, flexibility and stability. Nucleic acids research, 46(W1):W350–W355, 2018.

[35] Yuting Chen, Haoyu Lu, Ning Zhang, Zefeng Zhu, Shuqin Wang, and Minghui Li. Premps: Predicting the impact of missense mutations on protein stability. PLoS computational biology, 16(12):e1008543, 2020.

[36] Bian Li, Yucheng T Yang, John A Capra, and Mark B Gerstein. Predicting changes in protein thermodynamic stability upon point mutation with deep 3d convolutional neural networks. PLoS computational biology, 16(11):e1008291, 2020.

[37] S Benevenuta, C Pancotti, P Fariselli, G Birolo, and T Sanavia. An antisymmetric neural network to predict free energy changes in protein variants. Journal of Physics D: Applied Physics, 54(24):245403, 2021.

[38] Carlos HM Rodrigues, Douglas EV Pires, and David B Ascher. Dynamut2: Assessing changes in stability and flexibility upon single and multiple point missense mutations. Protein Science, 30(1):60–69, 2021.

[39] Yunzhuo Zhou, Qisheng Pan, Douglas EV Pires, Carlos HM Rodrigues, and David B Ascher. Ddmut: predicting effects of mutations on protein stability using deep learning. Nucleic Acids Research, page gkad472, 2023.

[40] Lasse M Blaabjerg, Maher M Kassem, Lydia L Good, Nicolas Jonsson, Matteo Cagiada, Kristoffer E Johansson, Wouter Boomsma, Amelie Stein, and Kresten Lindorff-Larsen. Rapid protein stability prediction using deep learning representations. Elife, 12:e82593, 2023.

[41] Corrado Pancotti, Silvia Benevenuta, Valeria Repetto, Giovanni Birolo, Emidio Capriotti, Tiziana Sanavia, and Piero Fariselli. A deep-learning sequence-based method to predict protein stability changes upon genetic variations. Genes, 12(6):911, 2021.

[42] Castrense Savojardo, Pier Luigi Martelli, Rita Casadio, and Piero Fariselli. On the critical review of five machine learning-based algorithms for predicting protein stability changes upon mutation. Briefings in bioinformatics, 22(1):601–603, 2021.

[43] Fabrizio Pucci, Katrien Bernaerts, Fabian Teheux, Dimitri Gilis, and Marianne Rooman. Symmetry principles in optimization problems: an application to protein stability prediction. IFAC-PapersOnLine, 48(1):458–463, 2015.

[44] Fabrizio Pucci, Katrien V Bernaerts, Jean Marc Kwasigroch, and Marianne Rooman. Quantification of biases in predictions of protein stability changes upon mutations. Bioinformatics, 34(21):3659–3665, 2018.

[45] Corrado Pancotti, Silvia Benevenuta, Giovanni Birolo, Virginia Alberini, Valeria Repetto, Tiziana Sanavia, Emidio Capriotti, and Piero Fariselli. Predicting protein stability changes upon single-point mutation: a thorough comparison of the available tools on a new dataset. Briefings in Bioinformatics, 23(2):bbab555, 2022.

[46] Dmitriy Umerenkov, Fedor Nikolaev, Tatiana I Shashkova, Pavel V Strashnov, Maria Sindeeva, Andrey Shevtsov, Nikita V Ivanisenko, and Olga L Kardymon. Prostata: a framework for protein stability assessment using transformers. Bioinformatics, 39(11):btad671, 2023.

[47] Felix Jung, Kevin Frey, David Zimmer, and Timo Mühlhaus. Deepstabp: A deep learning approach for the prediction of thermal protein stability. International Journal of Molecular Sciences, 24(8):7444, 2023.

[48] Yijie Zhang, Zhangyang Gao, Cheng Tan, and Stan Z Li. Efficiently predicting protein stability changes upon single-point mutation with large language models. arXiv preprint 2312.04019, 2023.

[49] Daniel J Diaz, Chengyue Gong, Jeffrey Ouyang-Zhang, James M Loy, Jordan Wells, David Yang, Andrew D Ellington, Alex Dimakis, and Adam R Klivans. Stability oracle: A structure-based graph-transformer for identifying stabilizing mutations. bioRxiv, pages 2023–05, 2023.

[50] Kit Sang Chu and Justin B Siegel. Protein stability prediction by fine-tuning a protein language model on a mega-scale dataset. bioRxiv, pages 2023–11, 2023.

[51] Henry Dieckhaus, Michael Brocidiacono, Nicholas Z Randolph, and Brian Kuhlman. Transfer learning to leverage larger datasets for improved prediction of protein stability changes. Proceedings of the National Academy of Sciences, 121(6):e2314853121, 2024.

[52] Rahul Nikam, A Kulandaisamy, K Harini, Divya Sharma, and M Michael Gromiha. Prothermdb: thermodynamic database for proteins and mutants revisited after 15 years. Nucleic acids research, 49(D1):D420–D424, 2021.

[53] Joicymara S Xavier, Thanh-Binh Nguyen, Malancha Karmarkar, Stephanie Portelli, Pâmela M Rezende, Jõao PL Velloso, David B Ascher, and Douglas EV Pires. Thermomutdb: a thermodynamic database for missense mutations. Nucleic acids research, 49(D1):D475–D479, 2021.

[54] Octav Caldararu, Rukmankesh Mehra, Tom L Blundell, and Kasper P Kepp. Systematic investigation of the data set dependency of protein stability predictors. Journal of Chemical Information and Modeling, 60(10):4772–4784, 2020.

[55] Jianwen Fang. A critical review of five machine learning-based algorithms for predicting protein stability changes upon mutation. Briefings in bioinformatics, 21(4):1285–1292, 2020.

[56] Fabrizio Pucci, Martin Schwersensky, and Marianne Rooman. Artificial intelligence challenges for predicting the impact of mutations on protein stability. Current opinion in structural biology, 72:161–168, 2022.

[57] Roshan M Rao, Jason Liu, Robert Verkuil, Joshua Meier, John Canny, Pieter Abbeel, Tom Sercu, and Alexander Rives. Msa transformer. In International Conference on Machine Learning, pages 8844–8856. PMLR, 2021.

[58] Zeming Lin, Halil Akin, Roshan Rao, Brian Hie, Zhongkai Zhu, Wenting Lu, Nikita Smetanin, Robert Verkuil, Ori Kabeli, Yaniv Shmueli, et al. Evolutionary-scale prediction of atomic-level protein structure with a language model. Science, 379(6637):1123–1130, 2023.

[59] Michael Heinzinger, Konstantin Weissenow, Joaquin Gomez Sanchez, Adrian Henkel, Martin Steinegger, and Burkhard Rost. Prostt5: Bilingual language model for protein sequence and structure. bioRxiv, pages 2023–07, 2023.

[60] Kotaro Tsuboyama, Justas Dauparas, Jonathan Chen, Elodie Laine, Yasser Mohseni Behbahani, Jonathan J Weinstein, Niall M Mangan, Sergey Ovchinnikov, and Gabriel J Rocklin. Mega-scale experimental analysis of protein folding stability in biology and design. Nature, 620(7973):434–444, 2023.

[61] Preethy Sasidharan Nair and Mauno Vihinen. V ari b ench: A benchmark database for variations. Human mutation, 34(1):42–49, 2013.

[62] Lijun Quan, Qiang Lv, and Yang Zhang. Strum: structure-based prediction of protein stability changes upon single-point mutation. Bioinformatics, 32(19):2936–2946, 2016.

[63] Stephen F Altschul, Thomas L Madden, Alejandro A Schäffer, Jinghui Zhang, Zheng Zhang, Webb Miller, and David J Lipman. Gapped blast and psi-blast: a new generation of protein database search programs. Nucleic acids research, 25(17):3389–3402, 1997.

[64] Michael Remmert, Andreas Biegert, Andreas Hauser, and Johannes Söding. Hhblits: lightning-fast iterative protein sequence searching by hmm-hmm alignment. Nature methods, 9(2):173–175, 2012.

[65] Milot Mirdita, Lars Von Den Driesch, Clovis Galiez, Maria J Martin, Johannes Söding, and Martin Steinegger. Uniclust databases of clustered and deeply annotated protein sequences and alignments. Nucleic acids research, 45(D1):D170–D176, 2017.

[66] Martin Steinegger, Markus Meier, Milot Mirdita, Harald Vöhringer, Stephan J Haunsberger, and Johannes Söding. Hh-suite3 for fast remote homology detection and deep protein annotation. BMC bioinformatics, 20:1–15, 2019.

[67] Thomas Wolf, Lysandre Debut, Victor Sanh, Julien Chaumond, Clement Delangue, Anthony Moi, Pierric Cistac, Tim Rault, Rémi Louf, Morgan Funtowicz, et al. Huggingface’s transformers: State-of-the-art natural language processing. arXiv preprint 1910.03771, 2019.

[68] Michel van Kempen, Stephanie S Kim, Charlotte Tumescheit, Milot Mirdita, Jeongjae Lee, Cameron LM Gilchrist, Johannes Söding, and Martin Steinegger. Fast and accurate protein structure search with fold-seek. Nature Biotechnology, pages 1–4, 2023.

[69] John Jumper, Richard Evans, Alexander Pritzel, Tim Green, Michael Figurnov, Olaf Ronneberger, Kathryn Tunyasuvunakool, Russ Bates, Augustin Žídek, Anna Potapenko, et al. Highly accurate protein structure prediction with alphafold. Nature, 596(7873):583–589, 2021.

[70] K Abdulla Bava, M Michael Gromiha, Hatsuho Uedaira, Koji Kitajima, and Akinori Sarai. Protherm, version 4.0: thermodynamic database for proteins and mutants. Nucleic acids research, 32(suppl 1):D120–D121, 2004.

[71] MD Shaji Kumar, K Abdulla Bava, M Michael Gromiha, Ponraj Prabakaran, Koji Kitajima, Hatsuho Uedaira, and Akinori Sarai. Protherm and pronit: thermodynamic databases for proteins and protein– nucleic acid interactions. Nucleic acids research, 34(suppl 1):D204–D206, 2006.

[72] Christine A Orengo, Alex D Michie, Susan Jones, David T Jones, Mark B Swindells, and Janet M Thornton. Cath–a hierarchic classification of protein domain structures. Structure, 5(8):1093–1109, 1997.

